# Impact of Direct Acting Antivirals on Survival in Patients with Chronic Hepatitis C and Hepatocellular Carcinoma

**DOI:** 10.1101/575670

**Authors:** William M. Kamp, Cortlandt M. Sellers, Stacey M. Stein, Joseph K. Lim, Hyun S. Kim

## Abstract

**Background:** To investigate the impact of direct-acting antivirals (DAA) and 12-week sustained viral response (SVR12) in patients with hepatocellular carcinoma (HCC) and chronic hepatitis C virus (HCV) infection.

**Methods:** Retrospective analysis of HCC patients diagnosed from 2005 to 2016 at an urban tertiary-care hospital. Kaplan-Meier curves and multivariable Cox proportional hazards models were used to assess survival.

**Results:** 969 patients met inclusion criteria. 478 patients received interventional oncology treatment (catheter-based therapies, ablation or combination locoregional therapies), 141 received supportive care (palliative or no treatment), 125 underwent liver transplantation, 112 had tumor resection and 94 received chemotherapy or radiation as their primary treatment. Median overall survival of the cohort was 24.2 months (95% CI: 20.9-27.9). 470 patients had HCV (56%). 123 patients received DAA therapies for HCV (26.2%), 83 of whom achieved SVR12 (68%). HCV-positive and HCV-negative patients had similar survival (20.7 months vs 17.4 months, p=0.22). Patients receiving DAA therapy had an overall survival of 71.8 months (CI: 39.5-not reached) vs 11.6 months (CI: 9.8-14.5) for patients without DAA therapy (p<0.0001). DAA patients who achieved SVR12 had an overall survival of 75.6 months (CI: 49.2-not reached) vs the non-SVR12 group (26.7 months, CI: 13.7-31.1, p<0.0001). Multivariable analysis revealed AJCC, Child-Pugh Score, MELD, tumor size, tumor location and treatment type had independent influence on survival (p<0.05). In HCV-positive patients, AJCC, MELD, tumor location, treatment allocation and DAA were significant (p<0.05). In patients receiving DAA therapy, only MELD and SVR12 were predictive of overall survival (p<0.05).

**Conclusions:** DAA therapy and achieving SVR12 is associated with increased overall survival in HCV patients with HCC.

**Summary:** Direct-acting antiviral use is associated with increased survival in hepatitis C-related hepatocellular carcinoma patients. Patients treated with direct-acting antiviral who achieved hepatitis C cure had additionally increased survival versus those treated with direct-acting antiviral who did not achieve hepatitis C cure. This study supports the use of direct-acting antiviral for hepatitis C treatment in hepatocellular carcinoma patients.

## Introduction

Hepatocellular carcinoma (HCC) represents a global public health burden, affecting an estimated 14 million persons worldwide, and is the third leading cause of cancer mortality.^1^ Within the United States, HCC is ranked 7^th^ for cancer related mortality and has seen a doubling in incidence from 1975 to 2007.^1,2,3,4^ The primary predisposing factors for HCC carcinogenesis is liver cirrhosis.^1^ Cirrhosis risk factors include chronic alcohol use, viral hepatitis, including Hepatitis C (HCV), and non-alcoholic fatty liver disease.^1^

Chronic HCV infection is the second most common risk factor for HCC and is responsible for 10-25% of all HCC cases.^1^ Over 20-30 years, 20-30% of patients with chronic HCV infections will develop cirrhosis and end stage liver disease and1-4% of these patients will progress to HCC each year.^5,6^ Of all HCV related HCC cases, 80-90% occur in the setting of cirrhosis.^2^ With more than 3.5 million patients in the United States and an estimated 130-170 million patients worldwide currently infected with HCV, the importance of HCV management in HCC therapeutic care and prevention is clear.^7,8^ The major current therapeutic goal for HCV and prevention of liver disease progression is sustained viral response (SVR), which is defined by negative HCV RNA at 12 weeks post-treatment (SVR12) and appears to be durable with a late virologic relapse rate of less than 1%.^9,10^

Therapeutic management of HCV has recently undergone a shifted from interferon-based therapies to all-oral interferon-free direct-acting antiviral (DAA) combination regimens. DAAs are a new class of drugs that target nonstructural proteins responsible for replication and infection of the hepatitis c virus.^10,11,12^ Genotype specific DAA therapies have been shown to reach SVR12 exceeding 90% of patients with fewer adverse effects compared with historic interferon-based regiments.^7,13,14,15,16,17,18^ SVR12 from DAA regimens have been associated with a decrease in liver outcomes including cirrhosis, hepatic decompensation, HCC and mortality.^19^ However, the impact of DAA regimens on clinical outcomes in patients with HCC remain limited. This study evaluates the impact of DAA on overall survival in HCV patients with HCC with the *a priori* hypothesis that SVR12 would be associated with improved outcome.

## Materials and Methods

### Study Cohort

Patient data were collected from those with prior consent to research participation and institutional review board approval. Patients met criteria if they had a radiologic or histopathologic HCC diagnosis defined by NCI/AASLD guidelines at Yale New Haven Hospital between 2005 to 2016. Patients were grouped by primary HCC treatment including liver transplantation, tumor resection, interventional oncological procedures (catheter-based therapies, ablation or combination locoregional therapy), systemic management (chemotherapy or radiation) and supportive care (palliative or no treatment). All patients were reviewed for history of HCV infection diagnosis based on positive HCV antibody, positive HCV RNA and/or ICD-9 recorded in electronic medical records. Only those with a HCV infection diagnosis were assessed for DAA treatment. Coinfections such as hepatitis B (HBV) and/or human immunodeficiency virus (HIV) or prior HCV treatment with therapies other than DAA and/or use of multiple DAAs did not preclude patients from analysis. SVR12 status was only collected for patients with HCV plus any reported DAA use for HCV. Patients that reached SVR12 on interferon-based regimens were not included with patients reaching SVR12 via DAA regimens. Exclusion criteria for this study included unknown survival status and histopathologic diagnosis of combined HCC and cholangiocarcinoma. Liver transplant patients were excluded from HCC, DAA and SVR12 overall survival and multivariable hazard ratio analyses. Patients were conservatively excluded in analyses for which they had unknown values. The primary outcome of interest was overall survival (OS), defined as time from HCC diagnosis to all-cause mortality or censoring.

### Statistical Analysis

Kaplan-Meier curves and Cox proportional hazards models were used to assess survival. Univariate analysis of age, sex, Child-Pugh Score, tumor size, model for end-stage liver disease (MELD), AJCC stage, body mass index, alpha-fetoprotein level, platelet count, unilobar or bilobar tumor presentation, presence of multiple tumors, main treatment, HCV infection, DAA treatment and SVR12 status were performed and variables with a p-value <0.2 were included into multivariable analysis. Statistical analyses performed with JMP Pro 13.1.0 (SAS Institute, Cary, North Carolina) and GraphPad Prism 8.0.0 (GraphPad Software, La Jolla, California). Values were considered statistically significant with a p-value of less than 0.05.

## Results

### Cohort Description

Between 2005 and 2016, 969 HCC patients met inclusion criteria (Table 1). Mean age of cohort at HCC diagnosis was 62.8±10.2 years. The group was predominately male at 79%. As shown in Figure 1A, 478 patients (49.3%) received interventional oncology therapies, 141 (14.6%) received supportive care, 125 (12.9%) underwent liver transplantation, 112 (11.6%) had tumor resection and 94 (9.7%) received chemotherapy and/or radiation as their primary treatment. Among non-transplant patients, 470 (57.0%) patients were HCV positive of which 123 (26.2%) received a DAA regimen. Of those patients receiving DAAs for HCV treatment 83 (67.4%) achieved SVR12 (Figure 1B). Median OS for all patients was 24.2 months (95% CI:20.9-27.9) (Figure 2A). Median OS for patients receiving liver transplantation (n=125) was not reached as more than 50% of subgroup were alive at time of last follow up. Patients undergoing tumor resection (n=112) had a median OS=56.7 months (95% CI: 41.9-103.5), interventional oncology (IO) (n=478) median OS=27.7 (95% CI:22.3-30.7), systemic therapy (n=94) median OS=5.6 months (95% CI: 4.4-7.3) and supportive management (n=141) median OS=2.4 months (95% CI: 1.9-3.2, overall p<0.0001, figure 2B).

**Table 1:** Cohort and Subgroup Characteristics. Characteristics of cohort, HCV, non-DAA, DAA, non-SVR12 and SVR12 subgroups, n (%) or mean ± standard deviation as marked. AJCC: American Joint Committee on Cancer stage, MELD: model for end stage liver disease, IO: interventional oncology, HCV: hepatitis C, DAA: direct-acting antivirals, SVR12: sustained viral response at 12 weeks.

|  | Cohort | HCV Patients | Non-DAA Patients | DAA Patients | Non-SVR12 Patients | SVR12 Patients |
| --- | --- | --- | --- | --- | --- | --- |
| Age at HCC Diagnosis (years) (mean±stdev) | 62.8±10.2 | 60.5±7.7 | 59.9±7.7 | 61.7±5.8 | 61.6±5.3 | 61.8±6.1 |
| Sex |  |  |  |  |  |  |
| Male | 768 (79.3%) | 390 (83.0%) | 202 (82.4%) | 99 (80.5%) | 28 (90.3%) | 63 (75.9%) |
| Female | 201 (20.7%) | 80 (17.0%) | 43 (17.6%) | 24 (19.5%) | 3 (9.7%) | 20 (24.1%) |
| Liver Factors |  |  |  |  |  |  |
| AJCC |  |  |  |  |  |  |
| 1 | 400 (43.8%) | 198 (44.5%) | 79 (34.5%) | 79 (65.3%) | 15 (48.4%) | 58 (71.6%) |
| 2 | 244 (26.7%) | 110 (24.7%) | 63 (27.5%) | 31 (25.6%) | 11 (35.5%) | 19 (23.5%) |
| 3 | 162 (17.7%) | 86 (19.3%) | 53 (23.1%) | 8 (6.6%) | 3 (9.7%) | 4 (4.9%) |
| 4 | 107 (11.7%) | 51 (11.5%) | 34 (14.8%) | 3 (2.5%) | 2 (6.5%) | 0 |
| Child Pugh Score |  |  |  |  |  |  |
| A | 512 (55.2%) | 254 (56.3%) | 116 (49.2%) | 84 (68.9%) | 24 (77.4%) | 54 (65.9%) |
| B | 284 (30.6%) | 132 (29.3%) | 80 (33.9%) | 30 (24.6%) | 6 (19.4%) | 21 (25.6%) |
| C | 132 (14.2%) | 65 (14.4%) | 40 (16.9%) | 8 (6.6%) | 1 (3.2%) | 7 (8.5%) |
| MELD (mean±stdev) | 11.4±5.7 | 10.7±4.6 | 11.4±5.2 | 9.5±3.4 | 9.3±3.1 | 9.67±3.5 |
| Tumor Size (cm) (mean±stdev) | 4.48±3.6 | 4.18±3.3 | 4.70±3.6 | 2.76±1.7 | 3.09±2.4 | 2.57±1.3 |
| Tumor Location |  |  |  |  |  |  |
| Unilobar | 628 (66.7%) | 298 (65.1%) | 137 (57.6%) | 93 (75.6%) | 17 (54.8%) | 68 (81.9%) |
| Bilobar | 314 (33.3%) | 160 (34.9%) | 101 (42.4%) | 30 (24.4%) | 14 (45.2%) | 15 (18.1%) |
| Multiple Tumors |  |  |  |  |  |  |
| yes | 442 (45.9%) | 225 (48.4%) | 133 (55.0%) | 47 (38.2%) | 19 (61.3%) | 27 (32.5%) |
| no | 520 (54.1%) | 240 (51.6%) | 109 (45.0%) | 76 (61.8%) | 12 (38.7%) | 56 (67.5%) |
| Main Treatment |  |  |  |  |  |  |
| Transplant | 125 (13.2%) | 0 | 0 | 0 | 0 | 0 |
| Resection | 112 (11.8%) | 56 (12.1%) | 21 (8.6%) | 21 (17.4%) | 5 (16.1%) | 16 (19.5%) |
| IO | 478 (50.3%) | 285 (61.6%) | 143 (58.4%) | 93 (76.9%) | 24 (77.4%) | 63 (76.8%) |
| Systemic | 94 (9.9%) | 45 (9.7%) | 26 (10.6%) | 4 (3.3%) | 1 (3.2%) | 1 (1.2%) |
| Supportive | 141 (14.8%) | 77 (16.6%) | 55 (22.4%) | 3 (2.5%) | 1 (3.2%) | 2 (2.4%) |
| Hepatitis C Factors |  |  |  |  |  |  |
| HCV Infection | 551 (56.9%) | 470 | 245 | 123 | 31 | 83 |
| DAA Treatment |  | 123 (26.2%) | 0 | 123 | 31 | 83 |
| SVR12 |  |  |  | 83 (67.5%) | 0 | 83 |

**Fig. 1:**
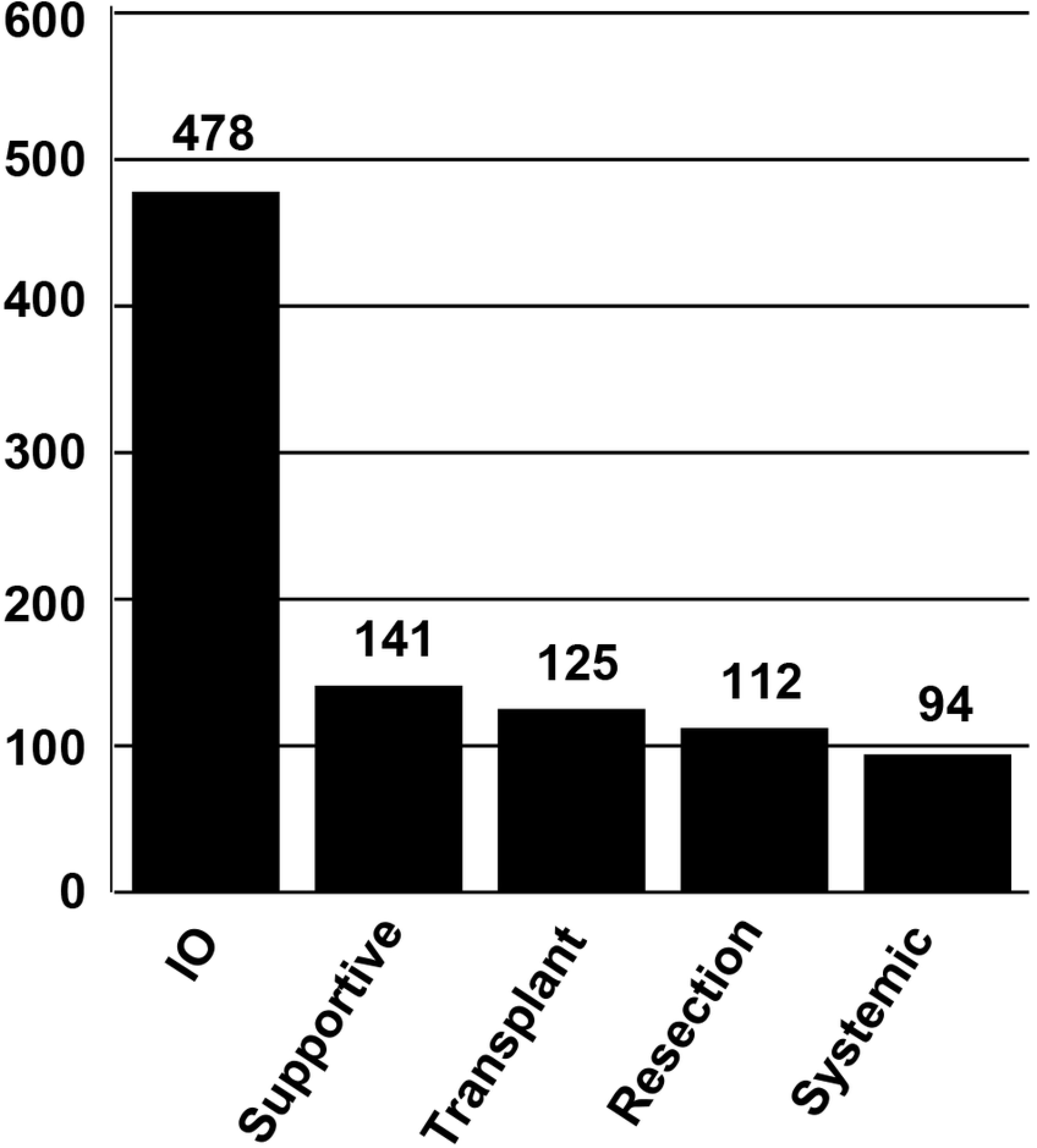
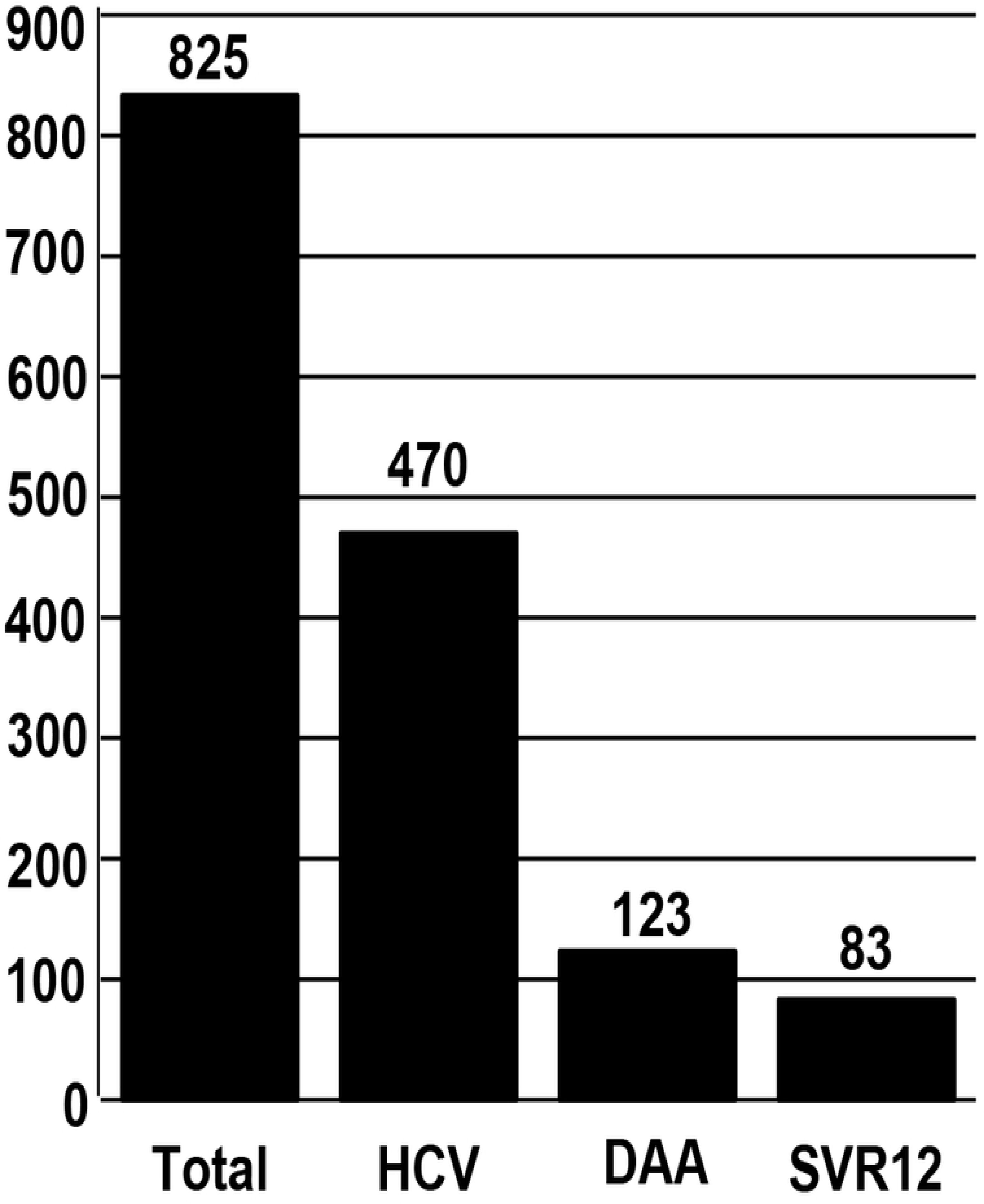
**A)** Treatment allocation of the entire hepatocellular carcinoma cohort **B)** Patients stratified by HCV infection, DAA therapy, and achievement of SVR12

**Fig. 2:**
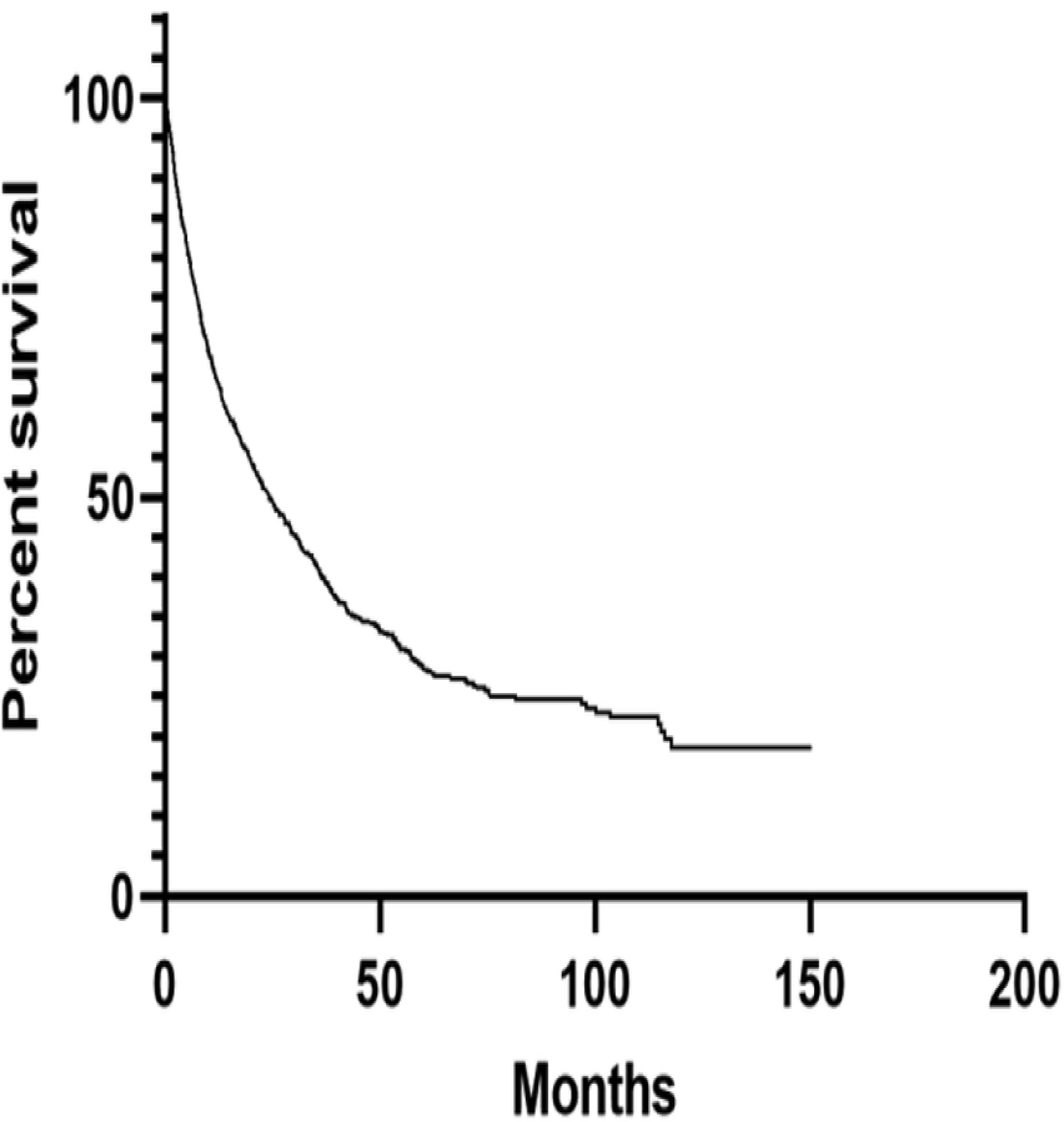
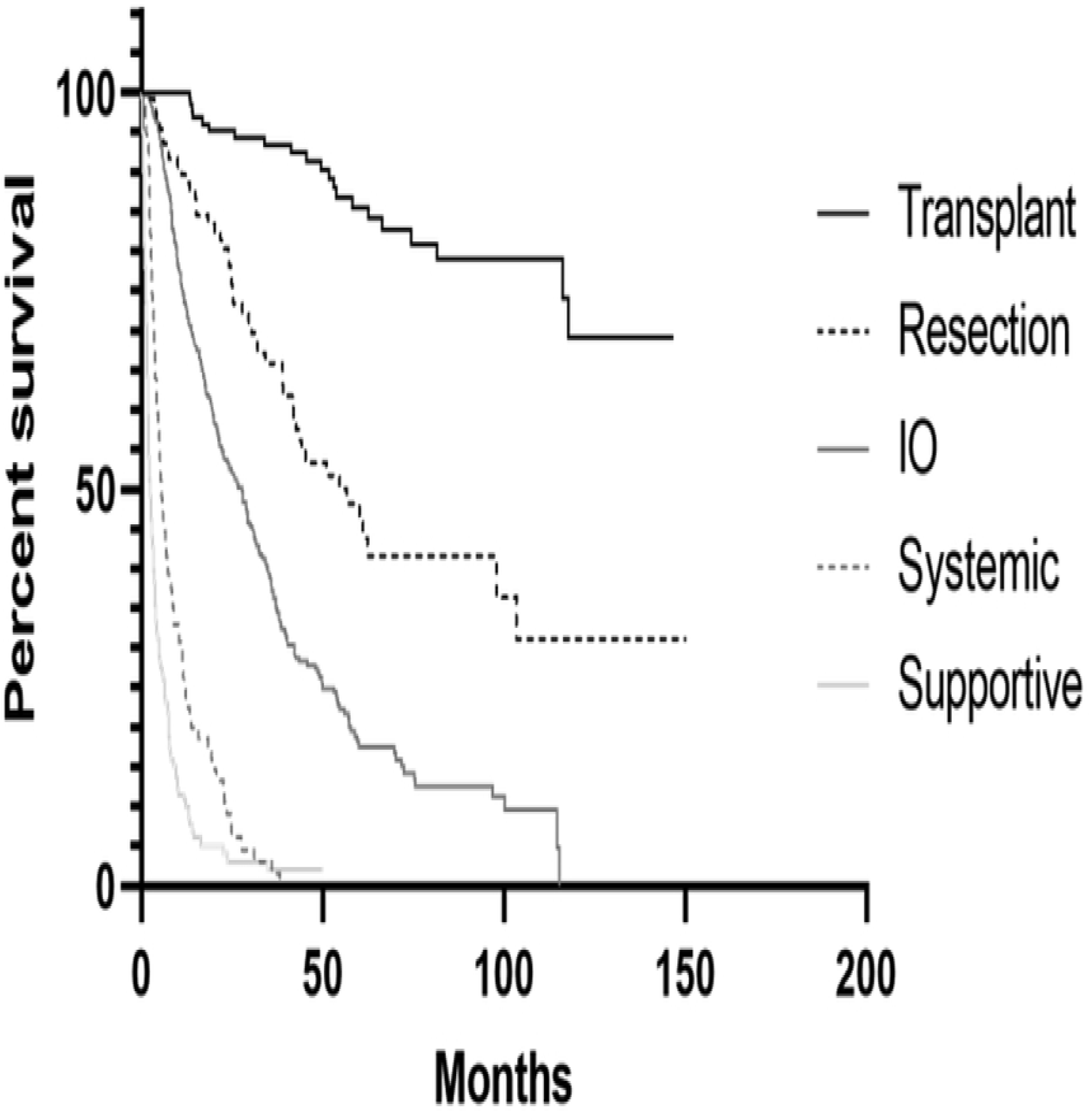
**A)** Survival rate for all hepatocellular carcinoma (HCC) patients within cohort (n=969) in months since HCC diagnosis. **B)** OS for HCC patients by main HCC treatment method. Patients receiving liver transplantation (n=125). Patients undergoing tumor resection (n=112), interventional oncology (IO) (n=478), systemic therapy (n=94) and supportive management (n=141) (overall p<0.0001).

### Overall Survival in HCV and DAA Subgroups

Subgroup analysis of HCV patients, recipients of DAA and those that achieved SVR12 revealed significant influences on OS. Although patients with and without HCV showed no significant difference in survival: median OS 20.7 months (95% CI: 16.5-24.1) versus 17.4 months (95% CI: 13.0-20.6) respectively, (p=0.22), HCV patients that received DAA had a median OS of 71.8 months (95% CI: 39.5-not reached) compared to 11.6 months (95% CI: 9.8-14.5) for HCV patients that did not use DAAs (p<0.0001)(Figure 3). Patients achieving SVR12 had a higher median OS of 75.6 months (95% CI: 49.2-not reached) versus 26.7 months (95% CI: 13.7-31.1) for patients with positive HCV RNA by PCR 12 weeks post-DAA cessation (Figure 3C) (p<0.0001).

**Fig. 3:**
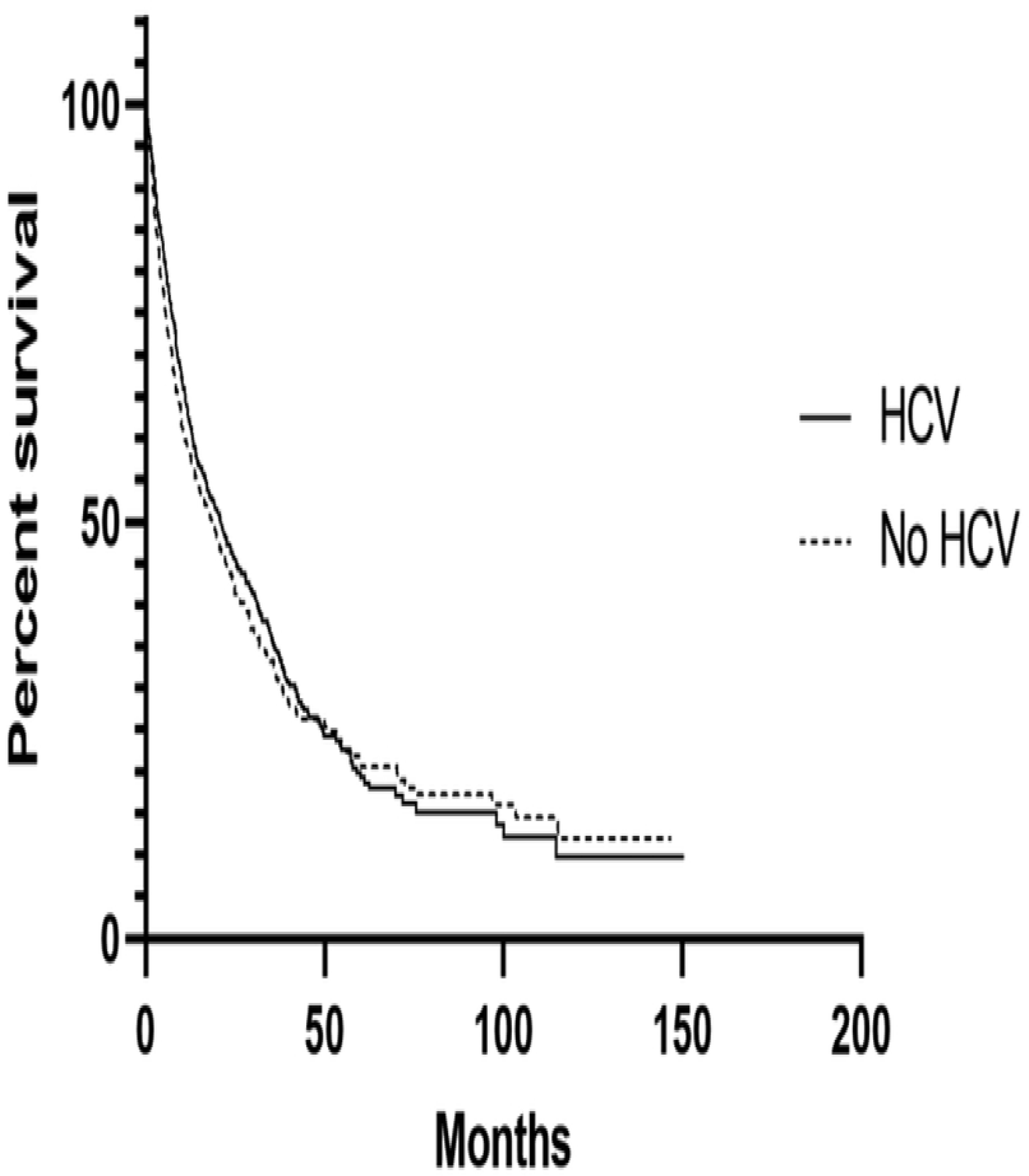
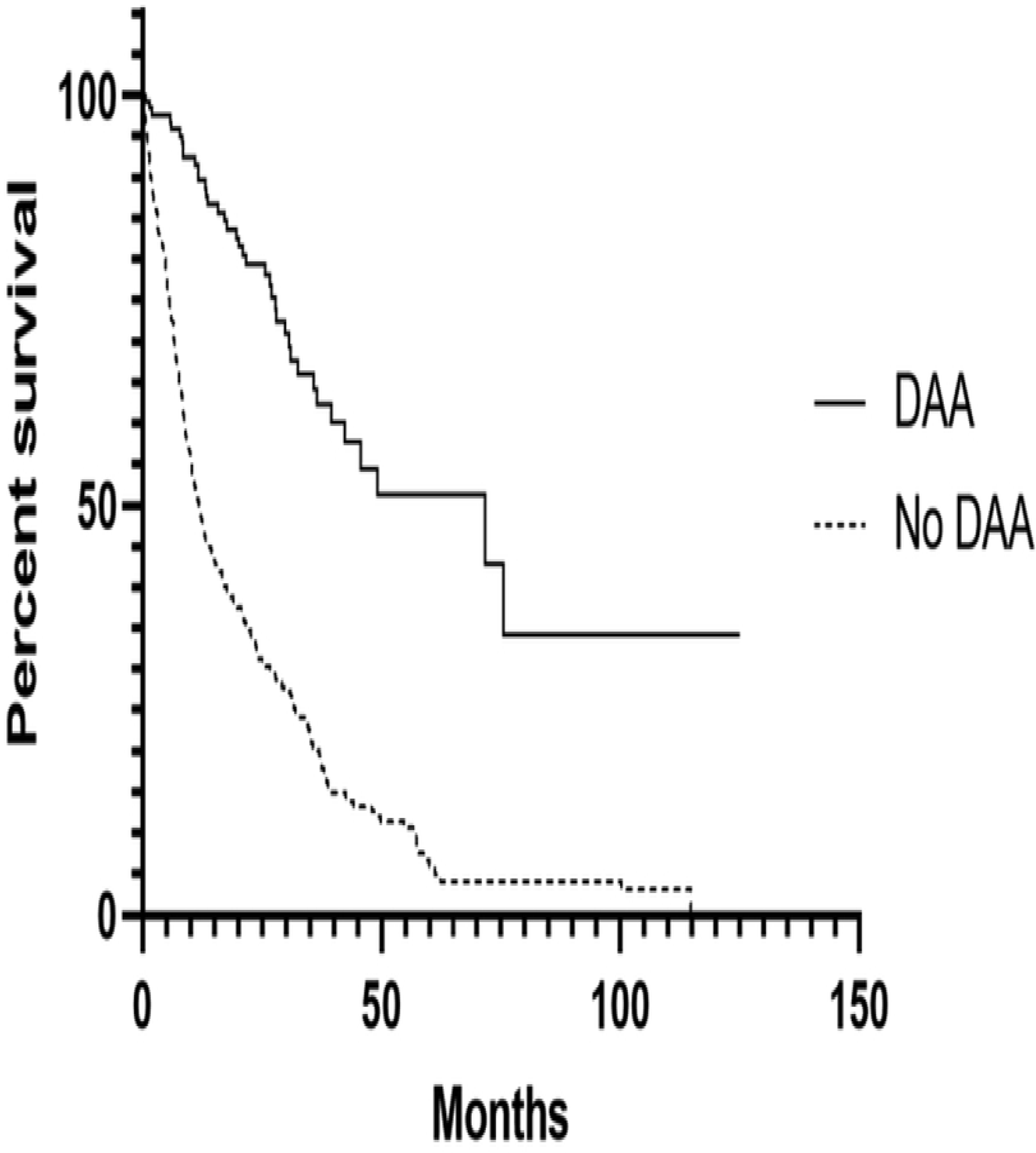
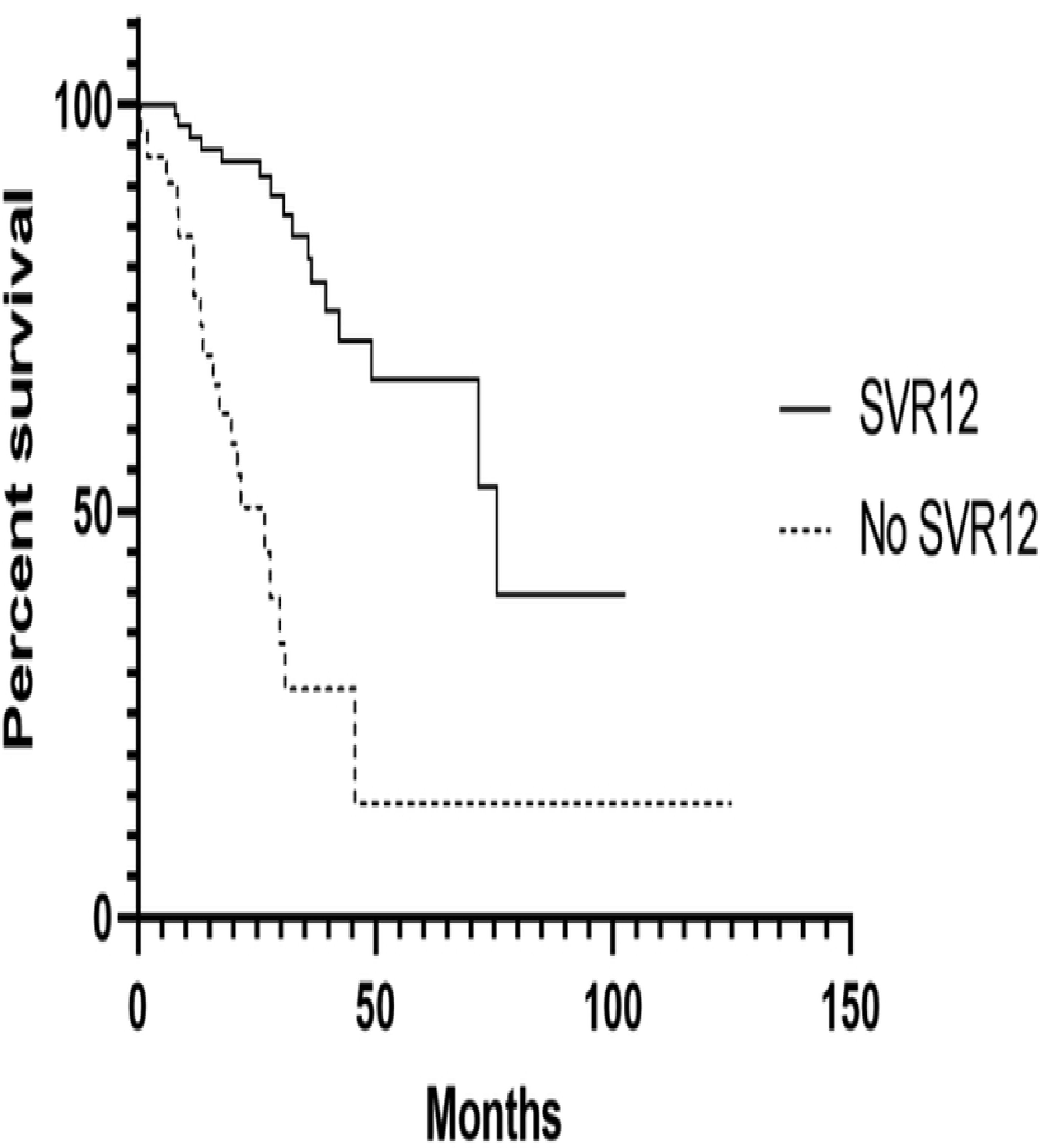
Patient survival rate in months since hepatocellular carcinoma (HCC) diagnosis. **A)** HCC Patients with positive history of hepatitis c (HCV) infection (n=470) versus patients with no history of HCV (n=363) (p=0.22) **B**) HCC and HCV patients that received a direct-acting antiviral (DAA) (n=123) versus those who did not receive a DAA (n=247) (p<0.0001) **C)** Patients with HCC and HCV that received a DAA and achieved sustained viral response (SVR12) (n=83) versus those who did not achieve SVR12 (n=31) (p<0.0001).

### Prognostic Factors

Multivariable analyses of subgroups were performed to assess impact of relevant HCC prognostic markers, HCC therapies, HCV factors and lab values. Within all non-transplant patients, numerous factors significantly influenced overall survival (Figure 4A, Table 2). AJCC stage 4 had decreased survival compared to stage 1 (HR=2.77, 95% CI:1.86-4.08, p<0.0001) and 2 (HR=2.06, 95% CI: 1.40-3.01, p=0.0003). AJCC stage 3 also had poorer survival as compared to stage 1 (HR=2.29, 95% CI: 1.26-2.30, p<0.0001) and 2 (HR=1.70, 95% CI: 1.26-2.30, p=0.0005). AJCC stage 2 had a HR=1.34 (95% CI: 1.03-1.75, p=0.03) compared to stage 1. Increased Child-Pugh score was also associated with worsened survival (Child Pugh C vs. A: HR 1.69, 95% CI: 1.07-2.65, p=0.02; B vs A: HR 1.83, 95% CI: 1.43-2.233, p<0.0001), as were increased MELD score, increased tumor size, and bilobar tumors (p<0.05). Treatment allocation significantly impacted survival, with resection demonstrating improved survival vs interventional oncology (HR=0.56, 95% CI: 0.39-0.81, p=0.002), systemic therapies (HR=0.26, 95% CI:0.16-0.42, p<0.0001) and supportive management (HR=0.12, 95% CI:0.08-0.19, p<0.0001). Interventional oncology increased survival rates over systemic therapies (HR=0.46, 95% CI:0.33-0.65, p<0.0001) and supportive management (HR=0.22, 95% CI:0.16-0.29, p<0.0001). Systemic therapies showed improved survival rates over supportive therapies (HR=0.47, 95% CI:0.42-0.68, p<0.0001).

**Table 2:** Hazard Ratios for Cohort and HCV and DAA subgroups. Hazard ratios from multivariable analysis. HR, 95% Confidence Interval, p-value, AJCC: American Joint Committee on Cancer stage, MELD: model for end stage liver disease, IO: interventional oncology, HCV: hepatitis C, DAA: direct-acting antivirals, SVR12: sustained viral response at 12 weeks.

|  | Total | HCV | DAA |
| --- | --- | --- | --- |
| Sex - Male/Female | 1.25 (0.99-1.49, p=0.0592) | 1.28 (0.89-1.89, p=0.1924) | 1.30 (0.39-5.25, p=0.6799) |
| Liver Factors |  |  |  |
| MELD | **1.03 (1.00-1.06, p=0.0168)** | **1.05 (1.00-1.11, p=0.0468)** | 1.21 (1.02-1.44, p=0.0326) |
| Tumor Size | **1.06 (1.02-1.09, p<0.0007)** | 1.01 (0.95-1.06, p=0.08082) | 0.75 (0.49-1.06, p=0.1044) |
| Bilobar/Unilobar | **1.32 (1.03-1.69, p=0.026)** | **1.62 (1.14-2.30, p=0.0072)** | 1.38 (0.36-5.51, p=0.6354) |
| Multiple Tumors - yes/no | 1.03 (0.80-1.32, p=0.816) | 1.27 (0.88-1.83, p=0.1986) | 1.01 (0.41-2.99, p=0.9831) |
| AJCC |  |  |  |
| 4/1 | **2.77 (1.86-4.08, p<0.0001)** | 1.59 (0.79-3.09, p=0.19) |  |
| 4/2 | **2.06 (1.40-3.01, p=0.0003)** | 1.32 (0.67-2.53, p=0.4209) |  |
| 4/3 | 1.21 (0.86-1.68, p=0.2704) | 0.75 (0.42-1.31, p=0.3186) |  |
| 3/1 | **2.29 (1.69-3.09, p<0.0001)** | **2.11 (1.25-5.54, p<0.0051)** | 1.81 (0.08-13.48, p=0.6350) |
| 3/2 | **1.70 (1.26-2.30, p=0.0005)** | **1.76 (1.05-2.92, p=0.0328)** | 2.28 (0.10-19.85, p=0.5336) |
| 2/1 | **1.34 (1.03-1.75, p=0.0305)** | 1.20 (0.79-1.82, p=0.3872) | 0.80 (0.27-2.22, p=0.6658) |
| Child-Pugh Score |  |  |  |
| C/A | **1.69 (1.07-2.65, p=0.0235)** | 0.86 (0.42-1.73, p=0.6775) | 0.13 (0.01-1.55, p=0.1068) |
| C/B | 0.93 (0.63-1.35, p=0.6904) | 0.65 (0.36-1.18, p=0.1549) | 0.15 (0.01-1.67, p=0.1234) |
| B/A | **1.83 (1.43-2.33, p<0.0001)** | 1.33 (0.92-1.91, p=0.1351) | 0.85 (0.30-2.40, p=0.7584) |
| Main Treatment |  |  |  |
| Resection/IO | **0.56 (0.39-0.81, p=0.0019)** | 0.79 (0.42-1.36, p=0.4118) | 0.39 (0.05-1.92, p=0.2649) |
| Resection/Systemic | **0.26 (0.16-0.42, p<0.0001)** | **0.29 (0.13-0.64, p=0.002)** |  |
| Resection/Supportive | **0.12 (0.08-0.19, p<0.0001)** | **0.12 (0.06-0.25, p<0.0001)** | 0.12 (0.00-5.53, p=0.2586) |
| IO/Systemic | **0.46 (0.33-0.65, p<0.0001)** | **0.37 (0.21-0.68, p=0.0017)** |  |
| IO/Supportive | **0.22 (0.16-0.29, p<0.0001)** | **0.16 (0.10-0.26, p<0.0001)** | 0.31 (0.01-9.82, p=0.4652) |
| Systemic/Supportive | **0.47 (0.32-0.68, p<0.0001)** | **2.34 (1.23-4.52, p=0.0093)** |  |
| Hepatits C Factors |  |  |  |
| HCV Infection - yes/no | 0.85 (0.70-1.04, p=0.1151) |  |  |
| DAA Therapy - yes/no |  | **0.39 (0.26-0.58, p<0.0001)** |  |
| SVR12 Attained - yes/no |  |  | **0.14 (0.06-0.35, p<0.0001)** |

**Fig. 4:**
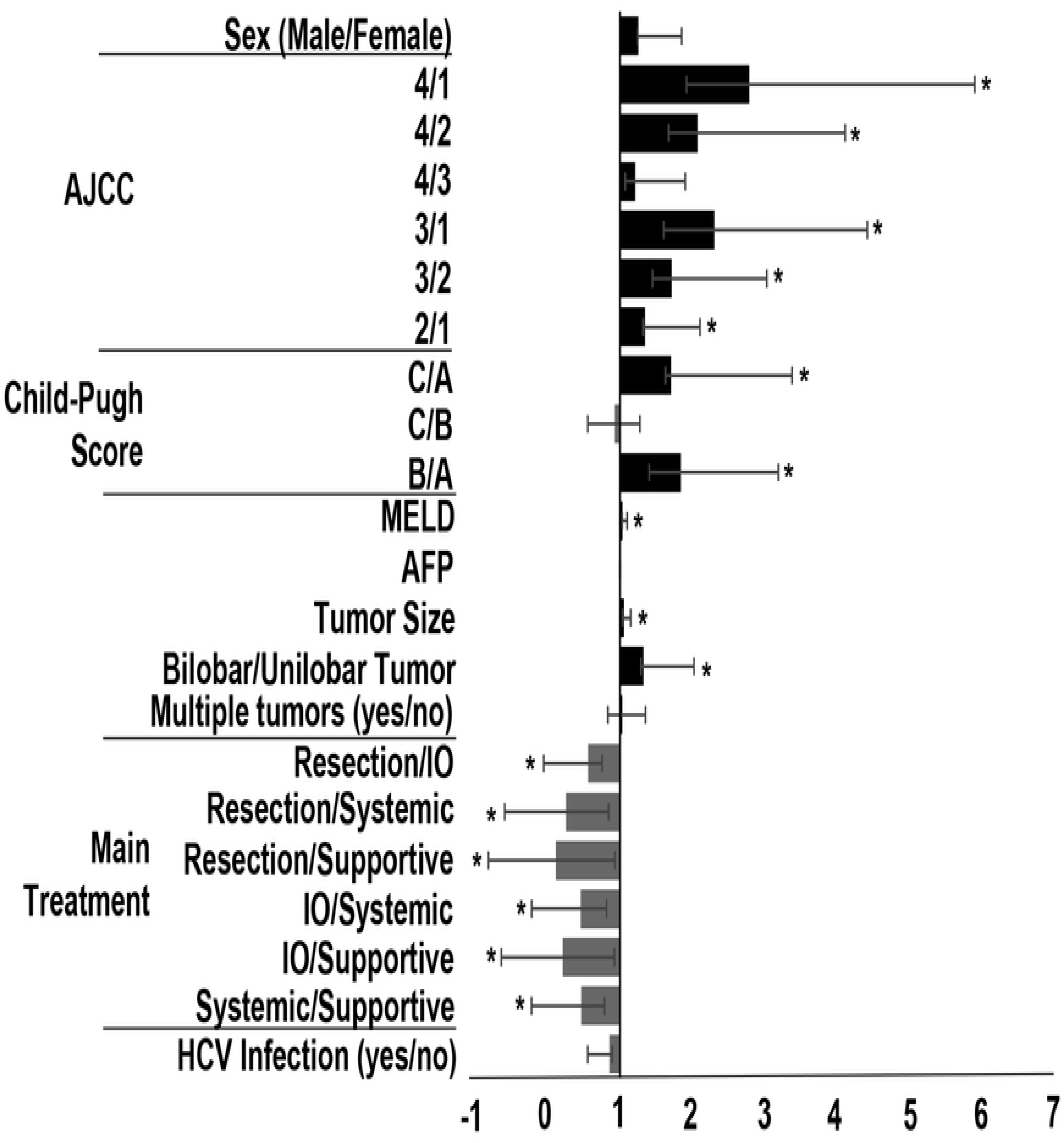
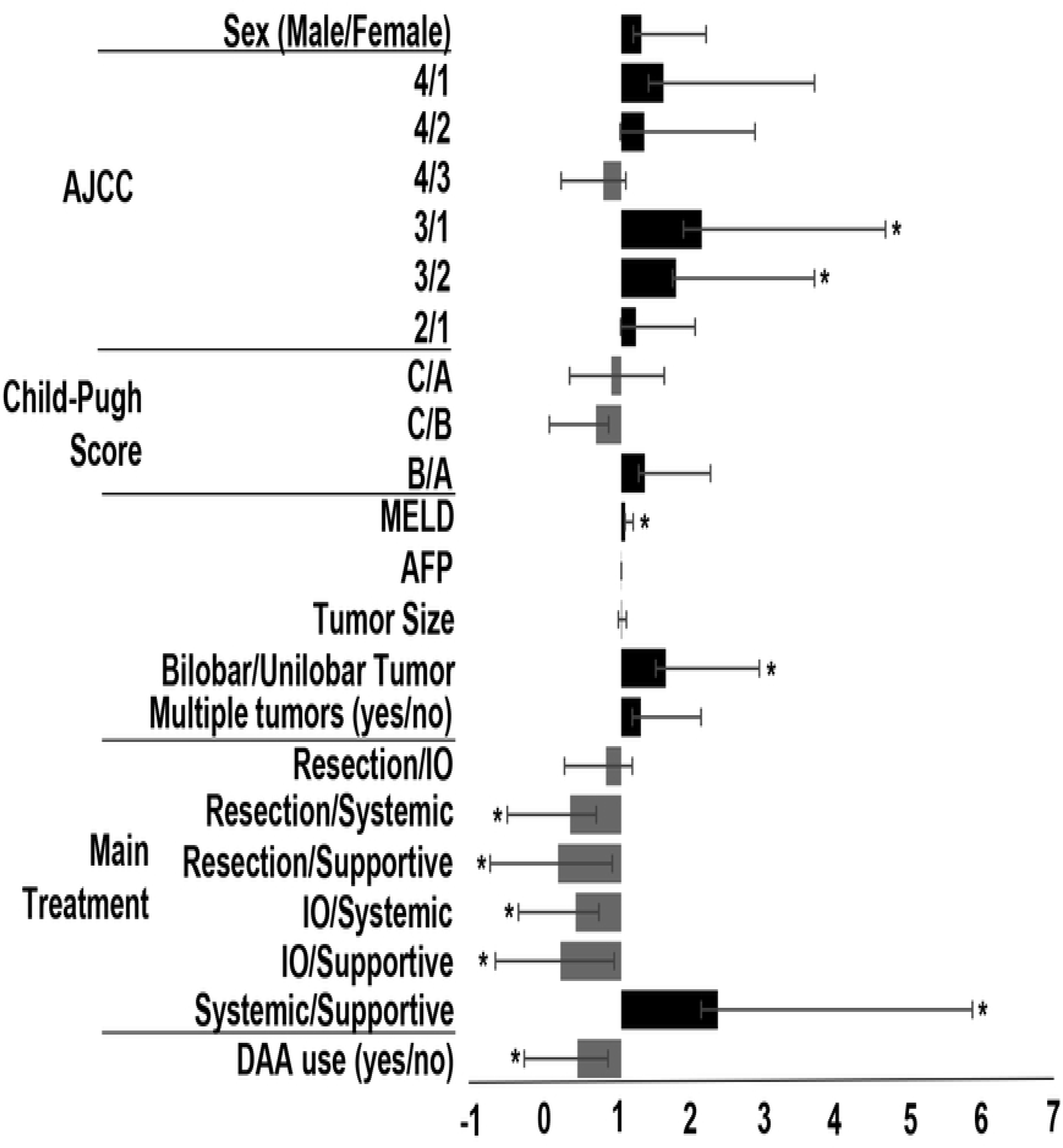
Hazard ratios from multivariable analysis on overall survival in non-transplant HCC patients. Values greater than one indicate increased risk of death. Values less than one indicate reduced risk of death. **A)** Hazard ratios in all non-transplant HCC patients. **B)** Hazard ratios in all non-transplant HCC with history of HCV. **\*** =p<0.05, AJCC: American Joint Committee on Cancer stage, MELD: model for end stage liver disease, AFP: alpha-fetoprotein, IO: interventional oncology, HCV: hepatitis C, DAA: direct-acting antivirals.

Multivariable analysis of non-transplant HCV patients showed similar results to non-transplant patients (Figure 4B) with the addition of DAA use as a significant prognostic marker (HR=0.39, 95% CI:0.26-0.58, p<0.0001). In the final subgroup, including only DAA patients, only two factors were significant, MELD (HR=1.21, 95% CI:1.02-1.44, p=0.03) and SVR12 (HR=0.14, 95% CI:0.06-0.35, p<0.0001).

## Discussion

With the increasing incidence and prevalence of HCC and the impact of ICV infection on development of cirrhosis and HCC tumor formation, the importance of understanding the effects of DAA in HCC patients is only becoming more vital. One-half of the HCC cases among the three-fold increase in HCC incidence between 1975 and 2007 in the US can be attributed to the aging chronic HCV population.^2^ Although there are indications that DAAs may slow progression to HCC,^20,21^ there remains a vast population of HCC patients that could potentially benefit from treatment of their chronic HCV infections.

It is likely that the improved median overall survival seen in our cohort among HCC patients taking DAAs is a direct result of the high success rate of achieving SVR12. Although more patients are needed to reduce the possible influence of DAA exclusion from patients with worse prognoses, the over three-fold difference in median overall survival between those that did and did not achieve SVR12 likely indicates profound longitudinal effects of HCV cure in HCC patients. Although our data indicates less severe HCC and liver disease in DAA and SVR12 patients versus their subgroup counterparts, multivariable analysis supports reduced all-cause mortality in patients receiving DAAs and achieving SVR12. In addition, others have reported that achieving SVR12 is associated with improved liver function, Child-Pugh scores and reversal of liver decompensation symptoms which could also be factors in the improved survival.^22,23^

Reaching SVR12 is not an easy task in the HCC patient population. Only 69% of our population reached SVR12 with similar results in other retrospective analyses of DAA use in HCC patients^24,25^ compared to reported values of over 90% in populations powered to DAA efficacy.^18^ Part of this may be due to the difficulty of integrating HCV treatment into HCC care, as demonstrated by the low rate of HCC patients with HCV receiving DAA treatment in our cohort (<50%). While there is currently much discussion as to how aggressive clinicians should be about treating active HCV in HCC patients, no official guidelines currently exist.^23,26^ Frequently, HCC management supersedes HCV treatment as many providers seek to triage HCV until after the cancer has been treated.^23^ In addition, debate is ongoing as to whether or not DAA use increases HCC recurrence rates.^18,20,27,28,29, 30^

Interestingly, our results suggest that there is ample time for HCV intervention in newly diagnosed HCC patients. The DAA treatment course typically 8-24 weeks,^31^ and patients receiving resection or IO treatments, 74% of our cohort population, had a median overall survival greater than 27 months. We are hopeful that conversion from non-DAA to DAA medications and subsequent SVR12 achievement will increase as awareness and use of DAA in current HCV management increases.

This cohort also raises important questions for future research. The patients achieving SVR12 with DAA include those that had previously failed interferon-based therapies and subsequently received DAA after their introduction, those that received multiple courses and combinations of DAA and those that received DAA for different lengths of time. Some received DAA during HCC management while others received DAA before their HCC diagnosis. More work is needed in areas concerning the differences in DAA regimen efficacy to understand the most appropriate combination of DAA and regimen length and then tailor these by HCV genotype, HCC prognosis and other host and genetic factors in order to help more patients achieve SVR12 and better overall outcomes.

## Author Contributions

William Kamp - Data collection and analysis, writing of article, editing of article, final approval of article

Cortlandt Sellers – Data collection and analysis, writing of article, editing of article, final approval of article

Stacey Stein – Writing of article, editing of article, final approval of article

Joseph Lim – Writing of article, editing of article, final approval of article

Hyun Kim – Concept and design, supervision, funding, data analysis, writing of article, editing of article, final approval of article

## Abbreviations

DAA: Direct-acting antivirals
SVR12: 12-week sustained virologic response
HCC: hepatocellular carcinoma
HCV: hepatitis c virus
OS: median overall survival
AJCC: American Joint Committee on Cancer stage
MELD: Model for end stage liver disease
HR: hazard ratio

